# The A-C Linker controls centriole cohesion and duplication

**DOI:** 10.1101/2024.10.04.616628

**Authors:** Lorène Bournonville, Marine. H. Laporte, Susanne Borgers, Paul Guichard, Virginie Hamel

## Abstract

Centrioles are evolutionarily conserved barrel-shaped organelles playing crucial roles in cell division and ciliogenesis. These functions are underpinned by specific structural sub-elements whose functions have been under investigation since many years. The A-C linker structure, connecting adjacent microtubule triplets in the proximal region, has remained unexplored due to its unknown composition. Here, using ultrastructure expansion microscopy, we characterized two recently identified A-C linker proteins, CCDC77 and WDR67, along with a newly discovered protein, MIIP. Our findings reveal that these proteins localize between microtubule triplets at the A-C linker, forming a complex. Depletion of A-C linker components disrupt microtubule triplet cohesion, leading to breakage at the proximal end. Co-removal of the A-C linker and the inner scaffold demonstrates their joint role in maintaining centriole architecture. Moreover, we uncover an unexpected function of the A-C linker in centriole duplication through torus regulation, underscoring the interplay between these protein modules.

## Introduction

Centrioles are evolutionarily conserved macromolecular complexes critical for a wide range of fundamental cellular processes, including cell division, motility, and signaling ^1,2^. In dividing cells, two centrioles are embedded within the pericentriolar matrix, together forming the centrosome, which acts as a microtubule-organizing center (MTOC) essential for the accurate segregation of genetic material into daughter cells ^3–5^. During ciliogenesis, centrioles, functioning as basal bodies, dock to the plasma membrane, initiating the formation of cilia or flagella—cellular extensions that serve as sensory antennas or drivers of fluid and cell movement ^6–9^.

Each sub-structural component of the centriole has a highly specific role in these biological processes. Human centrioles exhibit a nine-fold symmetrical, microtubule- based cylindrical structure, measuring approximately 450 nm in length and 250 nm in diameter ^10^. They are polarized along a proximal-distal axis, with distinct architectural features at each end. The distal extremity, about 50 nm long, contains microtubule doublets and is decorated with subdistal and distal appendages crucial for ciliogenesis^11,12^. The central core region spans approximately 250 nm where lies an inner scaffold, which is critical for maintaining centriole cohesion ^13–15^. Proximally, the cartwheel structure, approximately 150 nm long, is essential for centriole duplication and imparts the nine-fold symmetry of the organelle ^16–19^. The cartwheel is connected to the microtubule triplets (MTTs), consisting of a complete A-microtubule and incomplete B- and C-microtubules, via the pinhead structure ^10,20^. Neighboring proximal MTTs are bridged by the A-C linker, a structural element spanning 35-45% of the centriole length ^13,21^. Although its function remains enigmatic, it is hypothesized to contribute to the structural cohesion of the centriole ^22,23^. Additionally, the proximal region is encircled by an amorphous torus, composed in part of CEP152 and CEP63, which plays a key role in recruiting PLK4 ^24–26^. PLK4 phosphorylates STIL to initiate cartwheel formation through SAS-6 oligomerization, and procentriole assembly during the S phase, concurrent with DNA replication ^27–30^.

Despite advances in understanding the molecular composition and function of various centriole components, including the cartwheel, inner scaffold, and distal appendages ^13,31–41^, the A-C linker structure has remained poorly characterized since its initial description in the 1960s ^42^. The evolutionary conserved protein POC1 (Proteome Of Centriole 1) has been suggested as a component of the A-C linker, but its localization appears species-specific. For instance, in *Chlamydomonas reinhardtii*, POC1 is thought to localize to the proximal region of centrioles ^22,43^, while in human osteosarcoma cells, we have demonstrated that POC1B is a central core component ^13,14^. More recently, cryo-tomography analysis of POC1 in *Tetrahymena thermophila* proposed that POC1 is associated with the microtubule triplets, rather than the A-C linker, and is required for centriole structural integrity ^15^.

To elucidate the molecular identity of the A-C linker, we employed ultrastructure expansion microscopy (U-ExM), a technique that enables nanoscale protein mapping of centrioles ^44^. We previously screened 23 candidate proteins in mature centrioles from U2OS cells, and identified two poorly uncharacterized proteins, CCDC77 and WDR67, as putative A-C linker components ^21^.

Building on these findings, we aimed to unravel the function of the A-C linker. In addition to CCDC77 and WDR67, we identified a novel A-C linker component, MIIP (migration and invasion inhibitory protein). Using U-ExM in combination with cell biology approaches, we revealed two crucial functions of the A-C linker proteins —CCDC77, WDR67, and MIIP—in centriole cohesion but also in duplication through torus assembly. Collectively, this study establishes the role of the A-C linker in centriole function, offering a foundation for exploring its broader significance in cellular architecture and division processes.

## Results

### MIIP, together with CCDC77 and WDR67, localizes at A-C linker level in human centrioles

We identified recently CCDC77 and WDR67 as two putative A-C linker components ^21^. CCDC77 is a long coiled coil protein of 77KDa, and WDR67, also known as TBC1D31, is a 125 KDa protein containing 7 WD repeats, a 170 amino acid Rab-GAP-TBC domain, several coiled coil regions, and a C-terminal tail that mediates direct interaction with the E3 ubiquitin-protein ligase praja2 (PJA2) ^45^. We first capitalized on these two proteins to ask whether other proteins could be part of the A-C linker. To explore this, we conducted a cross DepMap analysis ^46,47^, which revealed co-dependencies between genes expression, specifically focusing on the relationships between CCDC77 and WDR67 (**Fig. 1a**). Among the top 100 genes, we identified 12 that shared co-dependencies with CCDC77 and WDR67: WDR8, SPICE1, CEP135, TUBE1, TUBD1, TEDC2, RTTN (rotatin), CEP44, CEP295, PP1R35, CEP152, and MIIP. Notably, 11 out of these 12 genes encode characterized centriolar proteins, most of which localize to the proximal end of the centriole. The exception is MIIP (Migration and Invasion Inhibitory Protein, also known as IIP45), a poorly characterized protein known for its role in inhibiting cell migration and invasion via its interaction with the insulin-like growth factor binding protein 2 (IGFBP-2) ^48^ (**Fig. 1a**). Although MIIP was previously detected in centrosome mass spectrometry analyses ^49^, it has not been characterized at the centriole level. Consequently, we decided to focus on MIIP for further investigation.

**Figure 1.**
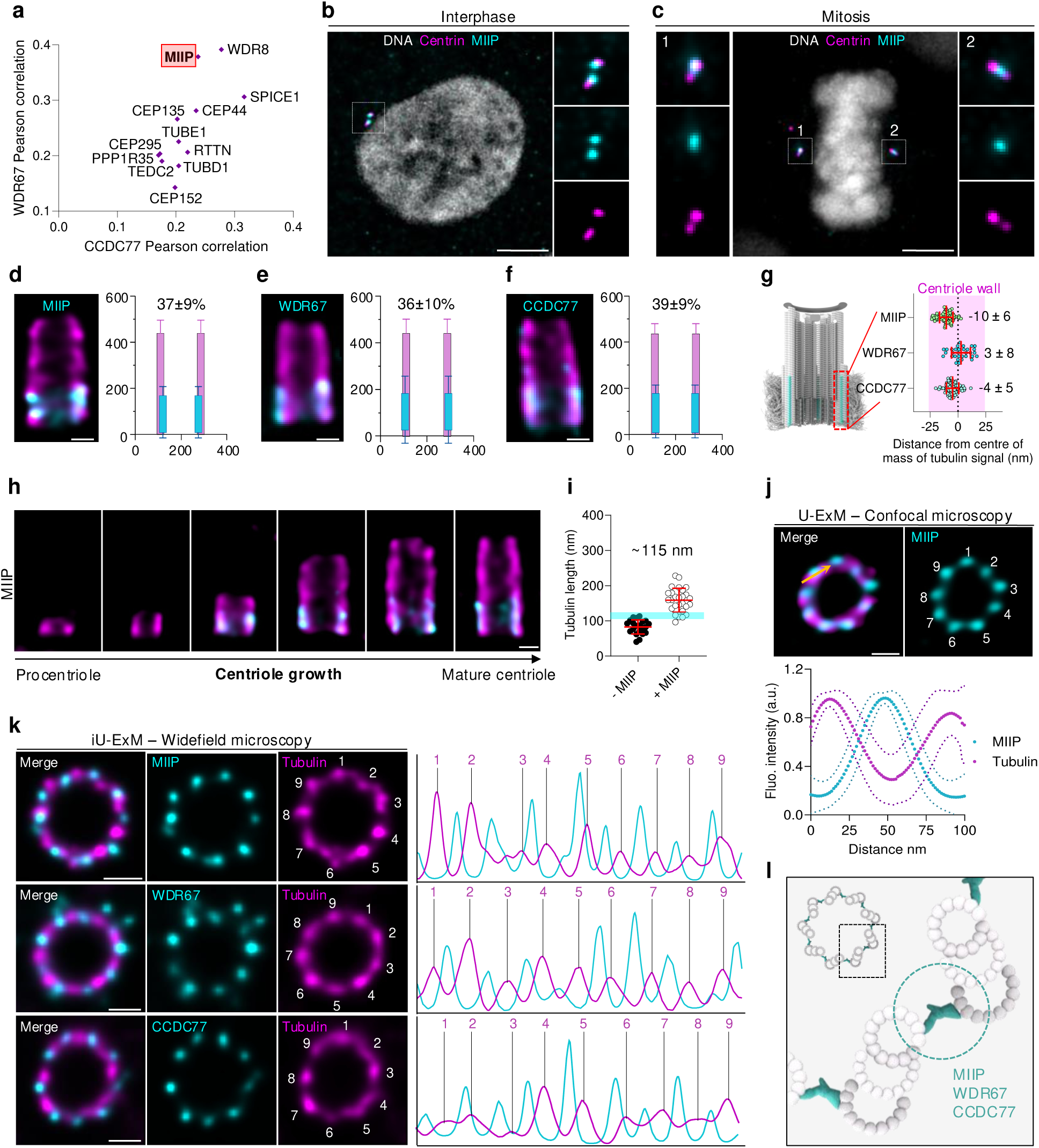
MIIP as a novel A-C linker protein **(a)** Common genes out of the top 100 genes plotted according to Pearson correlations of the WDR67 gene with the CCDC77 gene from the database Depmap portal (https://depmap.org/portal/). **(b-c)** Widefield images of U2OS cells in interphase (b) or mitosis (c) stained for DNA with DAPI (grey), Centrin (magenta), and MIIP (cyan). Scale bars = 5 μm. Dashed line squares correspond to insets. **(d-f)** Confocal images of expanded U2OS centrioles in longitudinal view stained for α/β-tubulin (magenta) and MIIP (cyan) (d), WDR67 (cyan) (e) or CCDC77 (cyan) (f). Scale bars = 100 nm. The average longitudinal and radial localization of MIIP (d) or WDR67 (e) or CCDC77 (f) are presented on the right of the corresponding image. **(g)** Model of a human centriole on the left displaying MTTs (gray) and the A-C linker structure (cyan) in the proximal part of the centriole. The relative radial positions of each protein of interest compared to tubulin are depicted next to the model. Error bars denote SD. **(h)** Confocal images of expanded U2OS procentrioles during assembly until the mature centriole stage in longitudinal view stained for α/β tubulin (magenta) and MIIP (cyan). Scale bar = 100 nm. **(i)** Centriole length, based on the tubulin signal, with (+ MIIP) and without (- MIIP) MIIP. The cyan line represents the average tubulin length when the MIIP signal appears (≈115 nm). **(j)** Confocal images of expanded U2OS centrioles from top view stained for α/β tubulin (magenta) and MIIP (cyan). Scale bar = 100 nm. MIIP signal is located between each MTT visualized thanks to tubulin staining. The fluorescence intensity profile along two successive MTTs demonstrating the precise position of MIIP is shown below the images. SDs are symbolized by the smaller dashed lines. The yellow arrow indicates the plot profile measurement of the MIIP signal between two successive MTTs. **(k)** Widefield images with Huygens deconvolution of expanded U2OS centrioles using iU-ExM from top view stained for α/β tubulin (magenta) and MIIP (cyan – top panel), WDR67 (cyan – middle panel) or CCDC77 (cyan – bottom panel). Scale bars = 100 nm. The nine MTTs are clearly visible with the tubulin signal. Fluorescence intensity profile of each protein (cyan) of interest through the walls (tubulin signal) of an entire centriole using the plugin “Polar Transform” from Fiji are presented next to the corresponding images. **(l)** Model of a human centriole in top view centered on the A-C linker region highlighted in cyan where CCDC77, WDR67 and MIIP localize. The detailed statistics of all the graphs shown in the figure are included in the Source Data file.

We first monitored the localization of MIIP at centrioles in U2OS cells using regular immunofluorescence microscopy. Our results confirmed that MIIP is consistently associated with centrioles throughout the cell cycle both in interphase **(Fig. 1b)** and mitosis (**Fig. 1c**). Using U-ExM to further gain in resolution and by staining for tubulin as a proxy for the centriolar microtubule wall, we found that MIIP localizes to the proximal region of centrioles and corresponds thus to be *a bona fide* centriolar component (**Fig. 1d and Extended data Fig. 1c**). Next, to further investigate whether MIIP could be a component of the A-C linker, we analyzed MIIP’s longitudinal localization in the proximal region and revealed that it spans 37% +/- 9 of the total centriole length, similarly to CCDC77 and WDR67’s coverages of 36% +/- 10 and 39% +/- 9 respectively (**Fig. 1d-f**) ^21^. The analysis of MIIP radial position relative to the microtubule wall showed that it localizes at the level of the MTT compared to the tubulin center of mass signal (-10 nm +/- 6), close to the values of CCDC77 (-4 nm +/- 5) and WDR67 (3 nm +/- 8) (**Fig. 1g**). Next using time series reconstructions of centriole assembly using tubulin as a proxy as we previously established ^21^, we monitored the appearance and elongation of MIIP. Like observed for CCDC77 and WDR67, we found that MIIP is recruited on procentrioles when the average tubulin length reaches around 115 nm (**Fig. 1h, i**), a length which marks the beginning of the elongation phase during centriole assembly ^21^.

Finally, to determine whether MIIP is positioned between microtubule triplets, where the A-C linker is expected to reside, we analyzed its distribution using U-ExM in top-viewed centrioles. A plot profile analysis revealed that MIIP forms 9 distinct foci, consistent with localization between the microtubule triplets, like the pattern seen with CCDC77 and WDR67 (**Fig. 1j**). However, due the 20 nm distance between microtubule triplets^20,50^ and the 60-70 nm resolution of U-ExM ^51^, we employed iterative expansion microscopy (iU-ExM) to achieve a higher resolution of 10 nm ^52^. This higher resolution confirmed that MIIP, along with CCDC77 and WDR67, is indeed localized between the microtubule triplets (**Fig. 1k**). Collectively, these findings demonstrate that MIIP is a novel component of the A-C linker in human cells (**Fig. 1l**)

### Co-dependency of CCDC77, WDR67 and MIIP complex

The A-C linker is a conserved centriole structural element whose complexity revealed in cryo-tomography suggests that it is composed by several proteins and that probably have multiple intricate interactions ^15,20,22,50^. Therefore, we wondered whether the loss of one of the three components identified in this study could lead to the destabilization of the others. To test this, we monitored the localization of CCDC77, WDR67 and MIIP in cells depleted of each of the individual components using siRNA treatment (**Fig. 2 and Extended data Fig. 1a-d**). We initially validated the specificity of each staining and the efficacy of each siRNA treatment by monitoring the overall centrosomal signals corresponding to CCDC77, WDR67, and MIIP in siCCDC77, siWDR67, and siMIIP conditions, respectively (**Fig. 2a-i**). Consistently with these proteins being structurally incorporated inside centrioles, we observed that the three proteins were mostly depleted from one centriole (53.4% in siCCDC77, 63.5% in siWDR67 and 51.7% in siMIIP), and more rarely in both centrioles (22.6% in siCCDC77, 29.5% in siWDR67 and 8.2% in siMIIP) (**Fig. 2e, g, i**). This result suggests that these proteins are stably incorporated and only the newly formed centriole is depleted. To discriminate between the mother and daughter centrioles, we co-stained cells depleted for CCDC77, WDR67 or MIIP with CEP164 (yellow arrow), a distal appendage protein present on mother centrioles ^53^ and the respective A-C linker proteins (**Extended Data** Fig. 1a-d). We found that the depletion occurred at the level of the daughter centrioles in the three conditions (**Extended Data** Fig. 1a-d), confirming that these three proteins are stably incorporated components of the centriole and mainly new centrioles assembled in the absence of the proteins are depleted.

**Figure 2.**
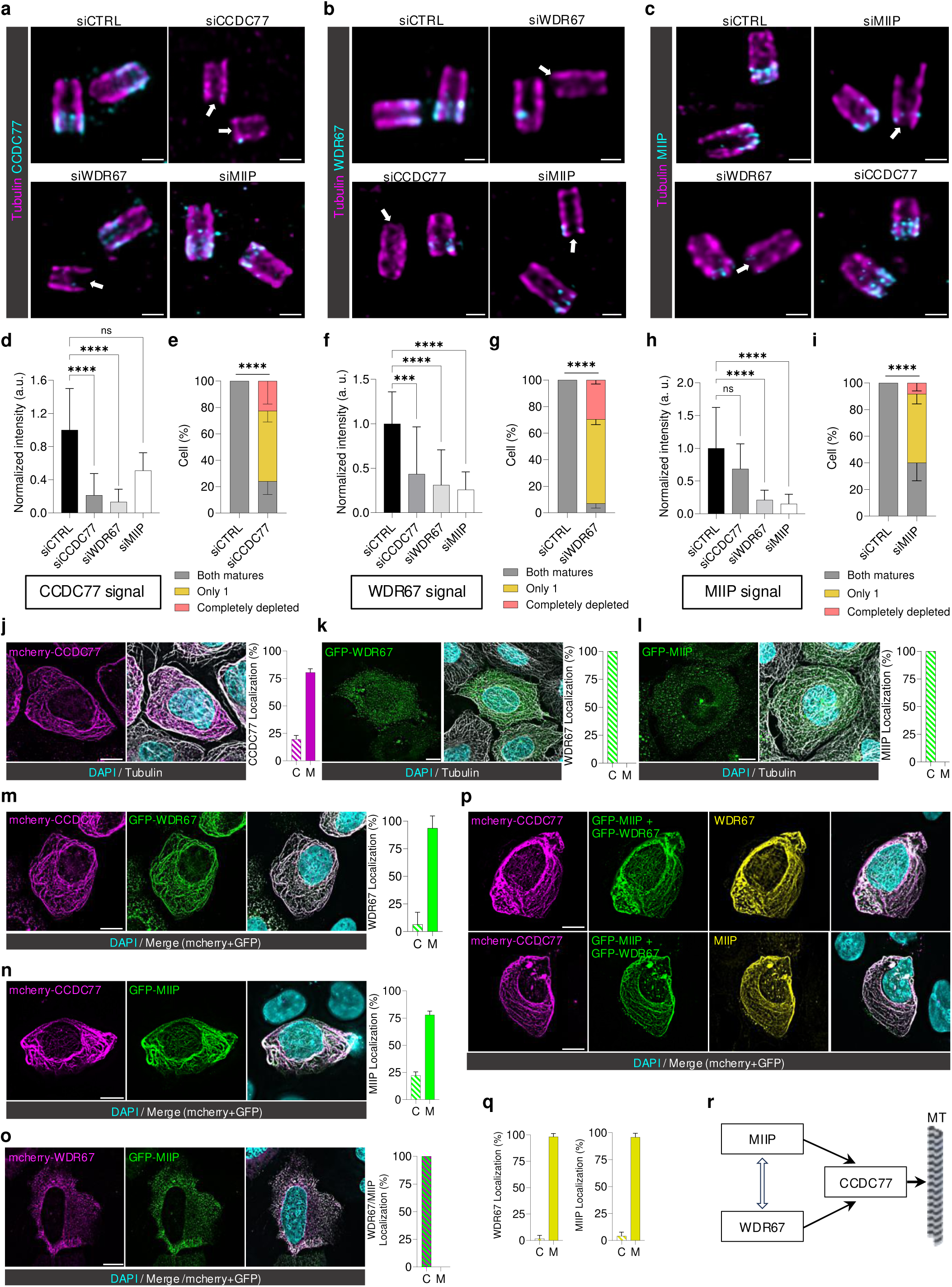
Co-dependency of CCDC77, WDR67 and MIIP complex (**a-c**) Confocal images of expanded U2OS centrioles in longitudinal view treated with siCTRL, siCCDC77, siWDR67, and siMIIP stained for α/β tubulin (magenta) and CCDC77 (a – cyan), WDR67 (b – cyan) or MIIP (c – cyan). Scale bars = 200 nm. White arrows point out the depletion of the protein targeted by each siRNA. (**d, f, h**) Normalized relative intensity of CCDC77 (d), WDR67 (f) or MIIP (h) in the different siRNA conditions. (**e, g, i**) Percentage of cells showing fluorescent signal for CCDC77 (e), WDR67 (g) or MIIP (i) in both mature centrioles (grey), only one (yellow) or none (orange) in the indicated siRNA (**j-l**) Widefield images of U2OS expressing mcherry- CCDC77 (j), GFP-WDR67 (k) or GFP-MIIP (l) were stained for DAPI (cyan) and α/β tubulin (grey). Scale bars = 10 μm. Percentage of cells showing CCDC77 (j), WDR67 (k) or MIIP (l) within the cytoplasm (C) or associated to microtubule (M) are presented next to the corresponding images. (**m-o**) Widefield images of U2OS cells co-expressing mcherry-CCDC77 (magenta) with GFP-WDR67 (green) (m), mcherry-CCDC77 (magenta) with GFP-MIIP (green) (n), mcherry-WDR67 (magenta) with MIIP-GFP (green) (o) stained for DAPI (cyan). Scale bars = 10 μm. Percentage of cells showing WDR67 (m), MIIP (n) or both (o) withing the cytoplasm (C) or associated to microtubule (M) are presented next to the corresponding images. (**p**) Widefield images of U2OS co-expressing mcherry-CCDC77 (magenta) with GFP-WDR67 (green) and GFP-MIIP (green) stained for DAPI (cyan) and WDR67 (yellow – top part) or MIIP (yellow – bottom part). Scale bars = 10 μm. (**q**) Percentage of cells showing WDR67 (left part) or MIIP (right part) within cytoplasm (C) or associated to microtubule (M). (**r**) Schematic view of the interactions between CCDC77, WDR67 and MIIP. MT stands for microtubule. The detailed statistics of all the graphs shown in the figure are included in the Source Data file.

Next, we monitored the impact of the depletion of the three A-C linker proteins on the localization of each other. Interestingly, we found that CCDC77 signal was strongly reduced upon siWDR67 but only mildly upon MIIP depletion (**Fig. 2a, d**).

Reciprocally, the fluorescent signal corresponding to WDR67 was strongly reduced upon siCCDC77 treatment (**Fig. 2b, f**). This result suggests that WDR67 and CCDC77 proteins are interdependent for their localization at the proximal region of centrioles. In contrast to CCDC77 however, we found that WDR67 localization was greatly impaired in siMIIP treatment (**Fig. 2b, f**), indicating that WDR67 localization at centrioles relies both on CCDC77 and MIIP. We next assess if MIIP localization was affected by CCDC77 or WDR67 depletion. Consistently, we found that MIIP distribution at centrioles was strongly reduced upon WDR67 depletion but only moderately upon CCDC77 depletion, probably as an indirect effect of the partial loss of WDR67 in that condition (**Fig. 2c, h**).

To further investigate whether the three identified A-C linker protein could interact and being part of the same complex, we used a microtubule displacement assay as previously published ^13,54^. We found that mcherry-CCDC77 binds microtubules (M) upon transient transfection in U2OS cells, in contrast to GFP-WDR67 and GFP-MIIP that stay cytoplasmic (C) or at the level of the centrosome (**Fig. 2j-l** and **Extended Data** Fig 1e). To assess whether CCDC77 can facilitate the displacement of WDR67 and MIIP onto microtubules, we co-transfected these proteins with mCherry-CCDC77. Remarkably, both GFP-WDR67 and GFP-MIIP were independently displaced onto microtubules (93% and 78%, respectively), indicating an interaction between these proteins (**Fig. 2m-o** and **Extended Data** Fig 1f-h). Notably, MIIP recruitment to microtubules was enhanced in the presence of WDR67 (**Fig. 2p, q**), suggesting that the three proteins can form a microtubule-associated complex (**Fig. 2r**).

### WDR67 and CCDC77 but not MIIP are important for ciliogenesis

We next investigated the function of the A-C linker components. CCDC77 and WDR67 were previously shown to be important for ciliogenesis ^45,55,56^, suggesting a potential role for the A-C linker in cilia formation. However, since both display additional localizations besides the A-C linker, at satellites ^55,56^ and at distal appendages for CCDC77 and at the proximal torus for WDR67 ^21^, it remains unclear whether their role in ciliogenesis is directly linked to the A-C linker. Given that MIIP is exclusively localized to the A-C linker, we investigated whether it also plays a role in ciliogenesis, to directly assess the involvement of the A-C linker structure in this process.

To do so, we treated RPE1 cells with either siRNA control or siCCDC77, siWDR67 or siMIIP and analyzed the impact on ciliogenesis by counting the number of cilia formed relative to the control (**Extended Data** Fig. 2). Consistently with previous reports ^45,55^, we found that 43% of CCDC77-depleted cells and 51% of WDR67-depleted cells displayed a primary cilium stained with acetylated tubulin, in contrast to the 73% observed in control cells, indicating that both proteins are important to regulate ciliogenesis (**Extended Data** Fig. 2a-e). Moreover, among the ciliated cells, 31% and 34% displayed shorter cilia in CCDC77-depleted and WDR67-depleted cells, respectively (**Extended Data** Fig. 2f, g). However, depletion of MIIP did not impact ciliogenesis (**Extended Data** Fig. 2d, e), suggesting that the ciliogenesis defect observed might be triggered by a mechanism independent of the A-C linker.

To further investigate the underlying causes of this phenotype and given that CCDC77 localizes to the distal appendages of mature centrioles, we used U-ExM to examine whether this localization was affected in cells depleted of CCDC77 or WDR67, while remaining intact upon MIIP depletion (**Extended Data** Fig. 2h-k). We observed that the distal localization of CCDC77 in RPE1 cells was significantly reduced upon depletion of CCDC77 and WDR67 but was less affected with MIIP (**Extended Data** Fig. 2h, i). A similar trend was noted in U2OS cells, where the reduction of CCDC77 at distal appendages was not significant upon MIIP knockdown (**Extended Data** Fig. 2j, k). These findings underscore a possible correlation between CCDC77 at appendages and ciliogenesis, suggesting that the ciliogenesis defects could be associated with the distal localization of CCDC77 rather than with the function of the A-C linker. In the same line, CCDC77 and WDR67 have been found at satellites, known to affect ciliogenesis ^55,56^. Altogether, these results indicate that the observed ciliogenesis defect is probably not due to A-C linker removal.

### The A-C linker maintains cohesion between the microtubube triplets in the proximal region

Since the A-C linker sub-element is bridging adjacent MTTs, a key prediction is that its loss would affect centriole architecture by destabilizing centriolar MTTs cohesion. To test this hypothesis, we analyze the centriole integrity using U-ExM in control and siRNA-mediated depletion of A-C linker components in U2OS cells (**Fig. 3**). Importantly, we found that the loss of CCDC77, WDR67 or MIIP resulted in approximately 25-30% of broken centrioles (**Fig. 3a-f**, white arrows). Additionally, re-expression of RNAi-resistant version of CCDC77, WDR67 or MIIP rescued this phenotype, highlighting the specificity of the observed effect (**Fig. 3a-f**).

**Figure 3.**
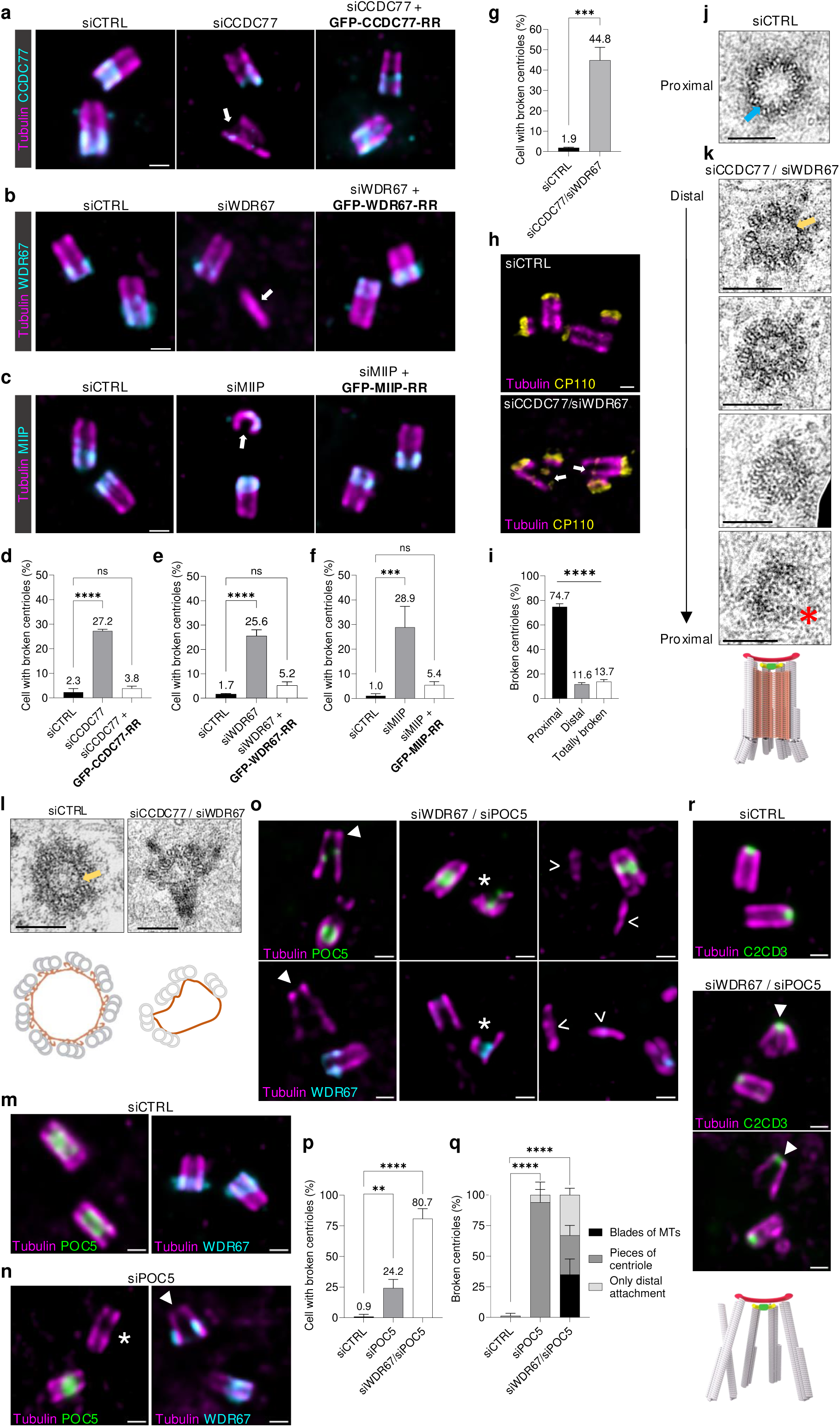
Depletion of A-C linker proteins leads to broken centriole (**a-c**) Widefield images of expanded U2OS stably expressing GFP alone or RNAi resistant GFP-CCDC77-RR (a), GFP-WDR67-RR (b) or GFP-MIIP-RR (c) stained for α/β tubulin (magenta) and the protein of interest (cyan) in different siRNA conditions. White arrows point to broken centrioles. Scale bars = 200 nm. (**d-f**) Percentage of cells with broken centrioles in the indicated siRNA conditions. **(g**) Percentage of cells with broken centrioles in the indicated siRNA conditions. (**h**) Confocal images of expanded U2OS cells treated with siCTRL or siCCDC77/siWDR67 and stained for α/β tubulin (magenta) or CP110 (yellow). White arrows point to broken centriole at the proximal region. Scale bar = 200 nm. (**i**) Percentage of cells with broken centrioles in different centriolar regions (proximal breakage or deformation, distal breakage or deformation or totally broken with only blades of microtubules or without any centriole shape). (**j**) Transmission electron microscopy images of U2OS centrioles treated with siCTRL. Scale bar = 200 nm. Blue arrow points to the A-C linker structure. (**k**) Transmission electron microscopy images of a U2OS centriole treated with siCCDC77/siWDR67 across the distal region to the proximal region. Scale bars = 200 nm. Yellow arrow points to the inner scaffold structure at the distal region. Red star points to a broken centriole at the proximal region. Human centriole model below indicates proximal breakage. (**l**) Transmission electron microscopy images of U2OS centrioles treated with siCTRL or siCCDC77/siWDR67. Yellow arrow points to the intact inner scaffold structure. Scale bars = 200 nm. Schematic views of the intact inner scaffold in siCTRL condition or the deformed inner scaffold/loss of MTTs in siCCDC77/siWDR67 are presented below to the corresponding images. (**m-o**) Widefield images of expanded U2OS centrioles treated with siCTRL (m), siPOC5 (n) or siWDR67/siPOC5 (o) stained for α/β tubulin (magenta) and POC5 (green) or WDR67 (cyan). Scale bars = 200 nm. White triangles point to broken centrioles with a preserved distal attachment; white stars point to pieces of broken centrioles; white arrowheads point to blades of microtubules from broken centrioles. (**p**) Percentage of cells with broken centrioles in the indicated siRNA conditions. (**q**) Percentage of cells with different types of broken centrioles with distal attachment or pieces of broken centrioles or blades of microtubules in the indicated siRNA conditions. (**r**) Widefield images of expanded U2OS centrioles treated with siCTRL or siWDR67/siPOC5 and stained for α/β tubulin (magenta) and C2CD3 (green). Scale bars = 200 nm. White triangles point to broken centrioles with a preserved distal attachment. Human centriole model below indicates a broken centriole with a distal attachment. The detailed statistics of all the graphs shown in the figure are included in the Source Data file.

We next wondered whether the dual depletion of A-C linker components would exacerbate the phenotype. To address this, we treated U2OS cells with siRNA targeting CCDC77 and WDR67 and analyzed their centriolar phenotype by U-ExM. We found around 45% of broken centrioles in this condition (**Fig. 3g, h**), confirming a synergistic effect of the co-depletion. We then sought to determine whether the weakening of the microtubule wall occurs along the entire length of the centriole or is more pronounced in the proximal region, where the A-C linker is localized. To investigate this, we stained U2OS control or CCDC77/WDR67-depleted cells with anti-tubulin antibodies to mark the microtubule wall, and the distal marker CP110 ^57^ (**Fig. 3h**). Analysis of the percentage of structural defects in the proximal, central or distal regions revealed that more than 75% of the centrioles with structural defects exhibited breakage primarily in the proximal region (**Fig. 3i**). These results indicate that the loss of A-C linker components particularly weakened the structural integrity of the proximal region of centrioles.

To further characterize this phenotype, we analyzed the ultrastructure of centrioles co-depleted for CCDC77 and WDR67 using resin-embedded electron microscopy. In control cells, centrioles displayed structurally intact and well-connected microtubule triplets with clear densities of the A-C linker (**Fig. 3j**). In contrast, serial section imaging of depleted cells revealed that the A-C linker densities are less visible at the proximal end, although the inner scaffold in the central region remained largely intact (**Fig. 3k**). In some cases, we observed central region centrioles with impaired structures, characterized by loss of MTTs and a deformed inner scaffold (**Fig. 3l**). These findings underscore the critical role of the A-C linker in maintaining the structural integrity and connectivity of MTTs.

Based on these results, we explored whether the observed centriole breakage could be an indirect consequence of the destabilization of other structural elements within the centriole. To address this, we examined the effects of depleting each A-C linker component on several other markers: CEP44, a protein located in the pinhead region in the proximal region of centrioles ^21,58,59^ (**Extended Data** Fig. 3a); CEP135 and SPICE, two proximal proteins ^21^(**Extended Data** Fig 3b, c); and POC5 for the inner scaffold ^60^(**Extended Data** Fig. 3d). We observed that depletion of A-C linker components resulted in mild impairments to all markers, with CEP44 and CEP135’s coverages being shorter and POC5 and SPICE’s coverage slightly extended (**Extended Data** Fig. 3a-h). To ensure no bias in the coverage quantification since we noticed a slight decrease in centriole size upon CCDC77 depletion (**Extended Data** Fig. 3i), we measured the length for each marker and confirmed our observations (**Extended Data** Fig. 3j-m). Overall, these results suggest that the depletion of A-C linker components subtly affects centriole architecture, indicating that the observed breakage is likely directly related to the loss of the A-C linker.

Finally, given that centrioles depleted for the A-C linker exhibit breakage predominantly in the proximal region while the inner scaffold remains intact, we investigated the impact of simultaneously removing these structures by co-depleting POC5, a component of the inner scaffold, and WDR67, an A-C linker component. Strikingly, 81% of centrioles were broken upon POC5 and WDR67 co-depletion in contrast to the 25-30% broken centrioles in the single depletion (**Fig. 3m-p**). This result demonstrates the complementary role of both the A-C linker and inner scaffold structures in maintaining MTT connection and centriole architecture integrity. Interestingly, we noticed that some impaired centrioles appeared to be maintained distally (**Fig. 3o, q**, white triangle). Staining with the distal marker C2CD3, which recently was shown to localize as an internal ring inside the centriole ^21,61^, confirmed this observation (**Fig. 3r**). This suggests the existence of a third structural connector located distally at the level of the microtubule doublet, which may contribute to the overall cohesion of the centriole along its proximal-distal axis.

### Depletion of A-C linker proteins impairs centriole duplication

Since the A-C linker is in the proximal region of centrioles and that centriole duplication arise from that region, we wondered whether the loss of the three A-C linker components could also impact centriole duplication. We first analyzed the number of centrin dots in mitosis as a marker of centriole duplication using immunofluorescence microscopy. We found that 52%, 59% and 54% of cells in mitosis had less than 4 centrin dots upon depletion of A-C linker components, indicative of a defect in centriole duplication (**Fig. 4a-i**). To ensure that the putative centriole duplication defects was real and not reflecting solely a reduction of centrin localization itself, we turned to U-ExM and monitored directly procentriole presence using tubulin as a proxy (**Fig. 4j-l**). We found that 24% of CCDC77 siRNA, 16% of WDR67 siRNA and 28% of MIIP siRNA-treated cells displayed 2 procentrioles in contrast to 50% observed in control cells (**Fig. 4j-o**), suggesting that the loss of these proteins impairs centriole duplication. Importantly, we verified that centriole duplication could be restored upon re-expression of an RNAi-resistant version of CCDC77, WDR67 and MIIP, ensuring the specificity of the described phenotype (**Fig. 4j-o**). Finally, we monitored the impact of the depletion on the cell cycle and found that the depletion of the A-C linker components was not impairing the cell cycle (**Extended Data** Fig. 4).

**Figure 4.**
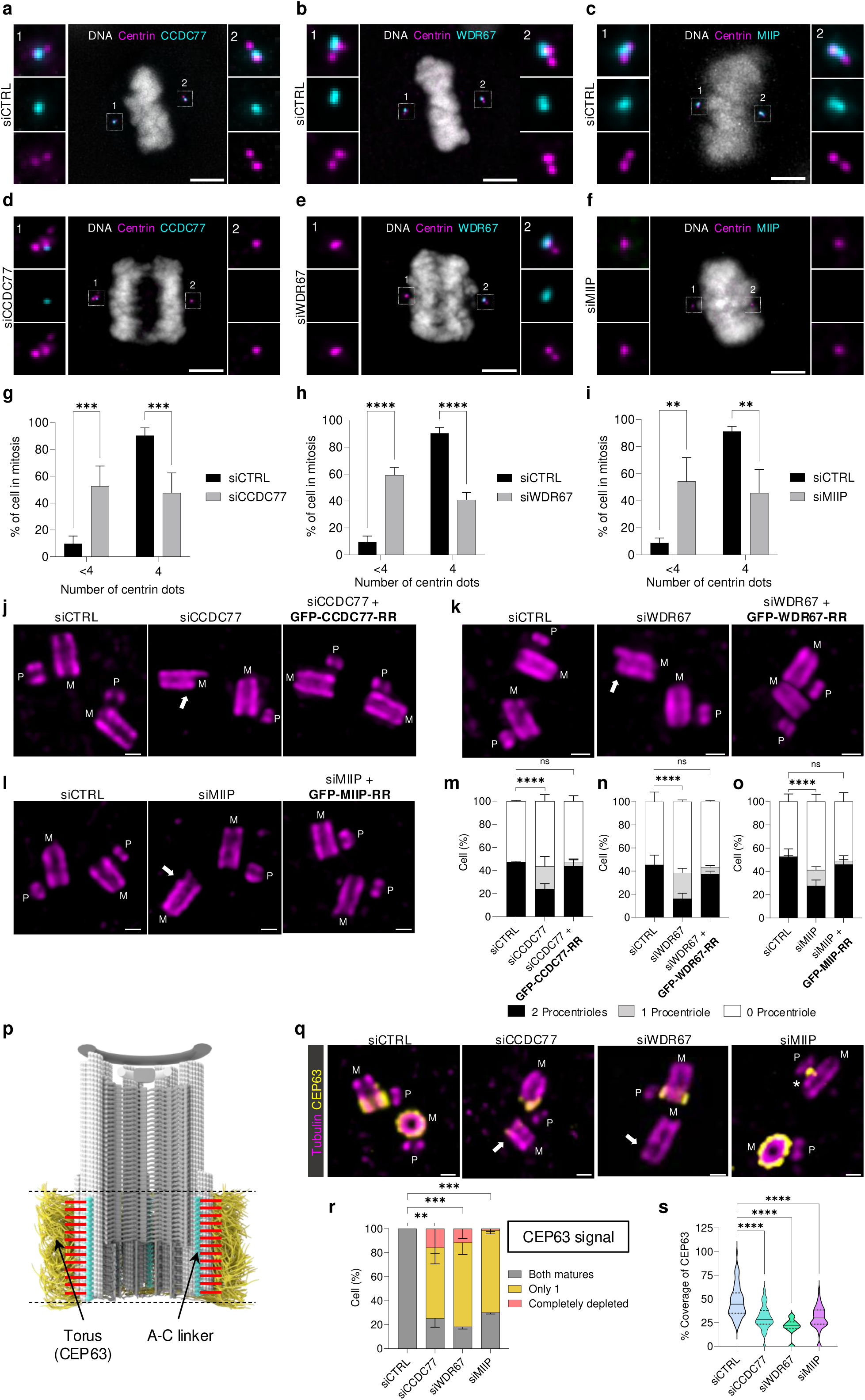
Depletion of A-C linker proteins impairs centriole duplication **(a-f)** Widefield images of U2OS cells in mitosis stained with DAPI (grey), Centrin (magenta), and CCDC77 (a, d – cyan), WDR67 (b, e – cyan) or MIIP (c, f – cyan) in different siRNA conditions. Scale bars = 5 μm. Dashed line squares correspond to insets. **(g-i)** Percentage of cell in mitosis with 4 or less than 4 centrioles (centrin dots) in different siRNA conditions. (**j-l**) Widefield images of expanded U2OS stably expressing GFP alone or RNAi resistant GFP-CCDC77-RR (j), GFP-WDR67-RR (k) or GFP-MIIP-RR (l) stained for α/β tubulin (magenta) and the protein of interest (cyan) in different siRNA conditions. Centrioles are in G2/S phase in longitudinal view. Scale bars = 200 nm. White arrows point to missing procentrioles. M stands for mature centriole and P stands for procentriole. (**m-o**) Percentage of cells with two procentrioles, only one or without any procentriole in the indicated siRNA conditions. (**p**) Model of a human mature centriole displaying the structural elements of the centriole and pointing to the A-C linker structure (cyan) and the torus (yellow) around the proximal region of the centriole. (**q**) Widefield images of expanded U2OS centrioles in G2/S phase stained for CEP63 (yellow) and α/β tubulin (magenta) in the indicated siRNA conditions. Scale bars = 200 nm. White arrows point to missing procentrioles. M stands for mature centriole and P stands for procentriole. Asterisk points to the remaining CEP63 signal at the base of the mother centriole. (**r**) Percentage of cells showing fluorescent signal for CEP63 in both mature centrioles (grey), only one (yellow) or none (orange) in the indicated siRNA conditions. (**s**) Coverage of CEP63 protein, expressed as a percentage of the tubulin length in the indicated siRNA conditions. The detailed statistics of all the graphs shown in the figure are included in the Source Data file.

**Figure 5.**
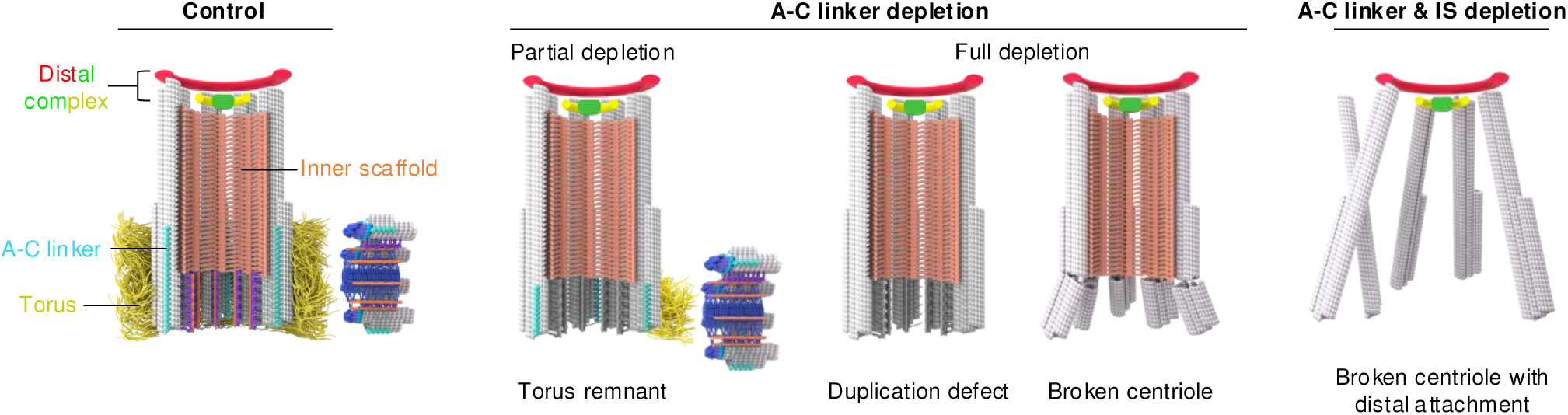
Functions of the A-C linker Schematic representation of a human centriole highlighting the distal complex, inner scaffold (IS), A-C linker and torus structural elements. This model underpins the functional roles of the A-C linker, made in parts of the proteins CCDC77, WDR67 and MIIP, in regulating centriole duplication through the torus localization, and maintaining centriole integrity.

Centriole duplication initiates around the torus, a fibrous density composed in part of CEP152 and CEP63, two proteins critical for recruiting proteins that initiate cartwheel and centriole formation ^59,62–64^. Since the A-C linker and the torus shares the same longitudinal position and length ^21^ (**Fig. 4p**), we next tested whether CEP63 localization would be affected upon CCDC77, WDR67 or MIIP depletion (**Fig. 4q, r**). We found that CEP63 signal was strongly reduced in the three tested conditions (**Fig. 4q-s**), unveiling an unexpected role for the A-C linker in dictating the recruitment of the CEP63 torus. Interestingly, we observed a correlation between procentriole presence and residual CEP63 signal at the base of the mother centriole (**Fig. 4q, asterisk**), indicating the robustness of centriole duplication, as even partial CEP63 signal appears sufficient to initiate procentriole formation. Collectively these results suggest that the A-C linker is not only crucial for maintaining the structural integrity of centrioles but also plays a pivotal role in ensuring the proper recruitment and localization of the torus, critical for centriole duplication mechanism.

## Discussion

The A-C linker, originally referred to as “A-C connections” by Gibbons and Grimstone in their pioneering 1960 study on flagellar structure in *Trichonympha, Pseudotrichonymph*a, and *Holomastigotes* ^42^, has since been widely recognized across various species, ranging from humans to *Paramecium* ^19,23,65–69^, with the exception of *Drosophila* ^70^ and the nematode *C. elegans*^41,71^. Despite these advances, the molecular composition of the A-C linker remained enigmatic. In this manuscript, we elucidate the identity and function of three key proteins—CCDC77, WDR67, and MIIP—as integral components of the A-C linker, revealing the functional significance of the A-C linker in human centrioles. We found that the A-C linker is crucial for maintaining the cohesion between microtubule triplets at the proximal end. We also found that the loss of the A-C linker slightly influences the other centriole’s structures. By simultaneously depleting the inner scaffold and A-C linker, we demonstrated that these two substructures together protect the centriole from fragmentation in an additive manner. Furthermore, we observed that depletion specifically affected only the daughter centriole. This suggests that the incorporation of the A-C linker is stable in mature centrioles, and only daughter centrioles assembled during the previous cycle under A-C linker depletion conditions lack it. This finding indicates that centriole assembly can occur even in the absence of the A-C linker. Finally, we also found that the A-C linker depletion leads to a centriole duplication defect due to the lack of the torus assembly in the newly formed centrioles lacking the A-C linkers. We conclude that the A-C linker has two critical functions at centriole level, microtubule cohesion and torus anchoring.

Previous study in cryo-electron tomography unraveled the high-resolution structure of the A-C linker in its native state in several species ^10,15,20,22,50,72,73^. These works demonstrated the complexity of this structural element that is composed of a A-link and a C-link, connected through a trunk ^22^. The whole structure is 15-20 nm long and is repeated every 8-8.5 nm along the microtubule triplets^22,50^. The molecular weight of WDR67 is around 124 KDa, 57KDa for CCDC77, and 43KDa for MIIP, for a total of 224 KDa. It is therefore possible that these three proteins constitute a large part of the A-C linker although it would be interesting in the future to determine whether other components are part of this sub-structural element. Our DepMap analysis identified other possible candidates, some already well characterized. It will be important later on to focus on these proteins in order to decipher the full molecular composition of the A-C linker. We have identified that CCDC77 associates with microtubules, suggesting that this protein is either involved in A-linker interaction with the A-microtubule or C- linker interaction with the C-microtubule. To determine its position, it will be necessary to develop super-resolution tools to pinpoint precisely which side the protein is on, or to increase the resolution of cryo-ET maps in order to place this protein unambiguously.

The role of the A-C linker in maintaining cohesion between the A and C microtubules of adjacent triplets has been proposed for some time. Our study confirms this role by identifying its components and demonstrating that the loss of the A-C linker results in breaks in the microtubule triplets at the proximal end. Unexpectedly, we discovered a second function of the A-C linker in torus anchoring. It has been well established that procentriole formation occurs proximally to the mature centriole. This duplication is enabled by the torus located in this region. Our observations revealed a positional correlation between the A-C linker and the torus. Removal of the A-C linker prevents torus recruitment, but intriguingly, partial depletion of the A-C linker is sufficient to recruit a reduced amount of torus and initiate procentriole duplication. These findings suggest that the A-C linker is crucial for defining the torus’s position at the proximal side of the mature centriole. Further studies are needed to investigate whether the displacement of A-C linker components affects procentriole duplication and to elucidate the significance of spatial positioning in centriole biogenesis.

## Methods

### Human cell lines and cell culture

Human bone osteosarcoma U2OS cells (ATCC-HTB-96) and retinal pigment epithelial cells hTERT RPE-1 (ATCC-CRL-4000) were grown in Dulbecco’s modified Eagle’s medium and GlutaMAX, supplemented with 10% fetal calf serum and penicillin and streptomycin (100 μg/ml) at 37 °C in a humidified 5% CO2 incubator. All cell cultures were regularly tested for mycoplasma contaminations.

To generate inducible episomal U2OS: GFP-CCDC77-RR cell line, U2OS cells were transfected using Lipofectamine 3000 (Life Technology). Transfected cells were selected or 6 days using 1 µg/mL puromycin starting day 2 after transfection. Selected cells were amplified and frozen. For further experiments, U2OS:GFP-CCDC77-RR, U2OS:GFP-WDR67-RR and U2OS:GFP-MIIP-RR cell lines were grown in the medium specified above supplemented with 1 µg/mL puromycin.

### Cloning

GFP-HsCCDC77-RR (RNAi resistant) was cloned in the Gateway compatible vector pEBTet-GFP-GW. An RNAi Resistant CCDC77 DNA sequence was synthesized by Geneart (ThermoFisher scientific) as such: position (636-653 bp) 5’ aCAcTAtCAgAGgGAtAT 3’ (modified region corresponding to siRNA from Thermo Fisher s38909) and position (1235-1252 bp) 5’ GcCGcATtCTcGAgGTtG 3’ (modified region corresponding to siRNA from Thermo Fisher s38908). The following restriction sites were modified: EcoRI and AgeI sites added at the 5’ end of the gene (note that the start codon has been removed), EcoRI (position 1127-1132 bp) mutated to 5’ GcATTC 3’, SalI (position 1232-1237 bp) mutated to 5’ GTCGcC 3’, XbaI and SalI sites were added at the 3’ end of the gene just after the stop codon. CCDC77-RR was first subcloned in pENTR using the restriction sites AgeI and XbaI. Subsequently, the final plasmid pEBTet-GFP-CCDC77-RR was obtained through a gateway reaction and sequence verified.

GFP-WDR67-RR (RNAi resistant) was cloned in the Gateway compatible vector pEBTet-GFP-GW. An RNAi resistant WDR67 DNA-sequence was synthesized by GeneArt (ThermoFisher scientific) as such: position (1411-1431 bp) 5’ aGaAAaCTgCTgAGgGTgTTa 3’ (modified region corresponding to siRNA from Thermo Fisher s228499) and position (2164-2187 bp) 5’ gAgGAcGAgGCcTGGTAtCAaAaa 3’ (modified region corresponding to siRNA from Thermo Fisher s228498). The following restriction sites were modified: AgeI site was added at the 5’ end of the gene (note that the start codon has been removed), and XbaI site was added at the 3’ end of the gene just before the stop codon. XbaI site (position 1840-1845 and 2905-2910) mutated to 5’ TCcAGg 3’. WDR67-RR was first subcloned in pENTR using the restriction sites AgeI and XbaI. Subsequently, the final plasmid pEBTet-GFP-WDR67-RR was obtained through a gateway reaction and sequence verified.

GFP-MIIP-RR (RNAi resistant) was cloned in the Gateway compatible vector pEBTet- GFP-GW. An RNAi resistant MIIP DNA-sequence was synthesized by GeneArt (ThermoFisher scientific) as such: position (111-131 bp) 5’- GAaTCgAGtCTaGAgTCtAgc-3’ (modified region corresponding to siRNA from Thermo Fisher s226949) and position (607-627 bp) 5’-cAaGAaTTcCGaGAgACtAAc- 3’ (modified region corresponding to siRNA from Thermo Fisher s34150). The following restriction site was modified: AgeI site (position 881-886) mutated to 5’ AtCGaTa 3’. MIIP-RR was first subcloned in pENTR using the restriction sites AgeI and XbaI. Subsequently, the final plasmid pEBTet-GFP-MIIP-RR was obtained through a gateway reaction and sequence verified.

### siRNA-mediated protein depletion and rescue experiments

U2OS cells were plated onto coverslips in a 6-well plate at 200.000 cells/well prior transfection and RPE1 cells were plated at 100.000 cells/well prior to transfection with the siRNA control and at 150.000 cells/well prior to transfection with les siRNAs against WDR67, CCDC77, and MIIP. Cells were next transfected 6h after with 20 nM silencer select negative control siRNA1 (4390843, Thermo Fisher) or siCCDC77 (s38908, sequence sense siCCDC77: 5’-GACGUAUCCUGGAAGUAGAtt-3’) or siWDR67 (s228498, sequence sense siWDR67: 5’- GAUGAAGCUUGGUACCAGAtt-3’) or siMIIP (s34150, sequence sense 5’- AGGAGUUUCGGGAAACCAAtt-3’), or siPOC5 (AD39Q91, sequence sense 5’- CAACAAAUUCUAGUCAUACUU-3’) using Lipofectamine RNAi MAX reagents (Invitrogen). Medium was changed 5-6 hours post-transfection. A second siRNA transfection was done 48h after the first one without changing the medium. Cells were analyzed 96 hours after the first transfection. Ciliogenesis was also performed under those conditions in RPE1 cells.

For the rescue experiments with U2OS:GFP-CCDC77-RR, or U2OS:GFP-WDR67- RR, or U2OS:GFP-MIIP-RR stable cell lines, the expression of the RNAi-resistant version of CCDC77 or WDR67 or MIIP was induced constantly for 96 hours using 1 µg/mL doxycycline.

### U-ExM protocol

The following reagents were used in U-ExM^74^ and iU-ExM^52^ experiments: formaldehyde (FA, 36.5-38%, F8775, SIGMA), acrylamide (AA, 40%, A4058, SIGMA), N, N-methylenbisacrylamide (BIS, 2%, M1533, SIGMA), sodium acrylate (SA, 97–99%, 408220, SIGMA and 7446-81-3, AK Scientific), ammonium persulfate (APS, 17874, ThermoFisher), tetramethylethylendiamine (TEMED, 17919, ThermoFisher), nuclease-free water (AM9937, Ambion-ThermoFisher), and poly-D-lysine (A3890401, Gibco). U2OS and RPE-1 cells were expanded using the U-ExM protocol as previously described (Gambarotto et al, 2019, 2021)). Briefly, cells were directly incubated for 3 hours in an anchoring solution containing 2% AA + 1.4% FA diluted in 1X PBS at 37 °C in a humid chamber. Next, the gelation step was performed using the U-ExM monomer solution (10% AA, 19% SA, and 0.1% BIS in 1X PBS) supplemented with 0.5% TEMED and APS by placing cells for 5 minutes on ice followed by 30 minutes at 37 °C incubation, followed by a denaturation step for 1 hours and half at 95 °C in a denaturation buffer (200 mM SDS, 200 mM NaCl, and 50 mM Tris in nuclease-free water, pH 9). Gels were washed from the denaturation buffer twice in ddH2O at room temperature for 30 minutes or overnight for complete expansion prior to immunostaining. Next, gels were measured with a caliper and the expansion factor was obtained by dividing the size after expansion by 12 mm, which corresponds to the size of the coverslips used for seeding the cells. For the immunolabelling, gels were placed in PBS for 15 minutes to shrunk and then incubated with primary antibodies in PBS-BSA 2% for 2 hours and half at 37°C. Next, 3 washes in PBS-Tween 0.1% were performed before to incubate the gels with secondary antibodies in PBS-BSA 2% for 2 hours at 37°C. Gels were washed 3 times in PBS-Tween 0.1% and then incubated 20 min at least in ddH20 for the final expansion.

### iU-ExM protocol

U2OS cells were expanded twice using the iU-ExM protocol as previously described^52^. Briefly, cells were directly incubated in the anchoring solution (2% AA; 1.4% FA in 1X PBS) for 3 hours at 37°C. Then, the gelation was performed using a homemade gelation chamber described in the iU-ExM protocol. The excess of the anchoring solution was removed using Kimwipes and the coverslip was glued on the slide of the gelation chamber. Next, this one was put on a humid chamber on ice and then a monomer solution (MS) (10% AA, 19% SA, 0.1% DHEBA, 0.25% TEMED/APS) was added to fill the space between the coverslip and the lid of the gelation chamber to completely cover the 12-mm coverslip. After 15 minutes on ice, the humid chamber was placed at 37°C for 45 minutes. This step was followed by the denaturation, the coverslip with the gel on top was carefully removed from the gelation chamber and dipped in 2 mL of denaturation buffer (200 mM SDS; 200 mM NaCl; 50 mM Tris-BASE; pH=6.8) in a 6-well plate under shaking until the gel detaches from the coverslip. Next, the gel was transferred in a 1.5 mL Eppendorf tube with 1 mL of fresh denaturation buffer and incubated for 1 hours and half at 85°C. A constant temperature is crucial for good gel consistency. After denaturation, several washes of ddH2O in a 12 cm petri dish were performed before a last wash overnight for a complete expansion. The intermediate antibody staining was done the next day as previously described for the U-ExM protocol after the first expansion step. Following the immunostaining step, the first expanded gel was cut to fit into a 6-well plate which was placed on ice. The piece of gel was incubated 25 minutes under shaking and on ice, with activated neutral gel (10% AA; 0.05% DHEBA; 0.1% APS/TEMED in ddH20). Then the gel was put on a microscope slide, and the excess of monomer solution was gently removed using kimwipes and it was covered by a 22 x 22 mm coverslip and incubated in a humid chamber for 1 hour at 37°C. Following this step, the gel embedded in the neutral gel was incubated in the anchoring solution (2% AA/1.4% FA) for 3 hours under shaking at 37°C. In a 6-well plate, the gel was washed in PBS 1X for 30 minutes and then incubated for 25 minutes under shaking and on ice with the 2^nd^ expansion monomer solution (10% AA, 19% SA, 0.1% BIS, 0.1% TEMED/APS) for a +/- 16X expansion factor. Next, the gel was transferred on a microscope slide, then the excess of monomer solution was gently removed with kimwipes and the gel was covered with a 22 x 22 mm coverslip for the incubation step 45 minutes at 37°C in a humid chamber. After final polymerization, the entire gel was incubated in 200 nM NaOH solution for 1h under agitation at room temperature in the dark for the dissolution of the first and neutral gels. Following this step, several washes with PBS (20 minutes in total) were performed before the final expansion in ddH2O, the water of which was changed several times until maximum expansion was reached after an overnight water bath.

### Imaging

Expanded gels were cut with a razor blade into squares to fit into a 36 mm metallic imaging chamber. The excess of water was carefully removed, and the gel was mounted onto 24 mm coverslips coated with poly-D-lysine (0.1 mg/mL) to prevent drifting. Images were taken with a 63x 1.4 NA oil immersion objective with either an inverted widefield Leica DMi8 Thunder microscope or a confocal Leica TCS SP8 microscope. For the widefield imaging, images were proceeded with the Thunder “Small volume computational clearing” mode and water as “Mounting medium” to generate deconvolved images. 3D stacks were acquired with 0.21 mm z-intervals and a 100 nm x, y pixel size. For the confocal imaging, images were proceeded with a lightning mode at max resolution, adaptative as “Strategy” and water as “Mounting medium” to generate deconvolved images. 3D stacks were acquired with 0.12 mm z-intervals and a 35-45 nm x, y pixel size.

### Antibodies used in this study

For immunostainings, primary antibodies used in this study were as follows: alpha-tubulin (AA345 scFv-F2C, abcd antibodies, 1:250), beta-tubulin (AA344 scFv-S11B, abcd antibodies 1:250), Acetylated tubulin (T7451, Merck Sigma-Aldrich, 1:500 for IF), CCDC77 (26369-1-AP, Proteintech, 1:500 for IF and 1:250 for U-ExM), WDR67 (HPA023710, Atlas antibodies, 1:500 for IF and 1:250 for U-ExM), MIIP (20630-1- AP, Proteintech, 1:500 for IF and 1:250 for U-ExM), Centrin (clone 20H5, 04-1624, Merck Millipore, 1:500 for IF), CEP63 (16268-1-AP, Proteintech, 1:500), CEP44 (24457-1-AP, Proteintech, 1:250), CP110 (12780-1-AP, Proteintech, 1:500), POC5 (A303-341A, Bethyl, 1:250), CEP135 (24428-1-AP, Proteintech, 1:250), C2CD3 (HPA038552, Atlas antibodies, 1:250), SPICE (A303-272A, Bethyl, 1:250), CEP164 (22227-1-AP, Proteintech, 1:250). Secondary fluorescent antibodies were purchased from Invitrogen ThermoFisher (anti-guinea pig 568 – A11075, anti-guinea pig 488 – A11073, anti-guinea pig Cy5 – Jackson Immuno Research 706-175-148, anti-rabbit 488 - A11008, anti-rabbit 568 – A11036, anti-rabbit 647 – A21245, anti-mouse 568 – A11004) and used at 1:800 dilutions for classical immunofluorescence and 1:400 for U-ExM).

### Immunofluorescence microscopy

U2OS or RPE-1 cells were grown on 12 mm coverslips and fixed at -20°C with cold methanol for 5 min. Fixed cells were then incubated with PBS-BSA 2% for 10 minutes at room temperature and next incubated with the primary antibodies for 1 hour at room temperature. Cells were subsequently washed three times with PBS-Tween 1% for 5 minutes and then incubated with the secondary antibodies conjugated with Alexa Fluor- 488 or 568. DNA was counterstained with DAPI solution. Samples were mounted in Fluoromount mounting medium (Fluoromount-G, 0100-01, SouthernBiotech) and observed with a widefield Leica DMi8 Thunder microscope. Images were taken with a 63x 1.4 NA oil immersion objective using the Thunder “Small volume computational clearing” mode and Fluoromount as “Mounting medium” to generate deconvolved images. 3D stacks were acquired with 0.21 um z-intervals and 100 nm x, y pixel size.

### EdU Cell proliferation Assay

Cells were treated for 96 hours with siRNAs and then with 10 μM EdU in complete medium for 30 minutes at 37°C. After incubation, cells were fixed at -20°C with cold methanol for 5 minutes and then washed twice with PBS-BSA 2%. The cells were next permeabilized with 0.5 Triton X-100 in PBS for 20 minutes at room temperature and again washed twice with PBS-BSA 2%. Detection of EdU incorporation into the DNA was performed with the Click-iT® EdU Alexa Fluor® 647 Cell Proliferation Assay Kit (Invitrogen, C10340) according to the manufacturer’s instructions. The Click-iT® EdU reaction cocktail (1×) was prepared according to the manufacturer’s instructions and added to the cell. Samples were incubated for 30 minutes at room temperature in the dark and then washed twice with PBS-BSA 2%. DNA was counterstained with DAPI solution. Samples were mounted in Fluoromount mounting medium and observed with a widefield Leica DMi8 Thunder microscope. Images were taken with a 20x 0.4 NA objective using the Thunder “Small volume computational clearing” mode and Fluoromount as “Mounting medium” to generate deconvolved images.

### Displacement assay

U2OS cells were plated onto coverslips in a 6-well plate at 300.000 cells/well prior transfection with different plasmid: mcherry-CCDC77 (gift from Juliette Azimzadeh’s laboratory), mcherry-WDR67 (gift from Juliette Azimzadeh’s laboratory), pEBTet- GFP-WDR67-RR and pEBTet-GFP-MIIP-RR. Cells were transfected the next day with 2.5 μg DNA per well using jetPRIME transfection reagents (Polyplus). Medium supplemented with doxycycline (1 µg/mL) was changed 5-6 hours post-transfection and cells were analyzed 24 hours post-transfection by classical immunofluorescence microscopy.

### Quantification

#### Data representations and quantifications

Images were analyzed using Fiji^75^. Only raw data were used for intensity measurements, otherwise only deconvolved images were used for representations and quantifications. Measurements of the length, the relative position, and the coverage of centriolar proteins were done as previously described^21^. Briefly, images were resized by decreasing the pixel size by 6 to improve the precision by using the plugin “CropAndResize”. The fluorescent signal distribution of tubulin and the protein of interest were measured using the Fiji line scan and the plugin “PickCentrioleDim” to easily select the start and the end of the fluorescent signal defined as 50% of the peak value at both extremities of the centriole. For all the measures, the tubulin was defined as the reference protein, and its starting coordinate was shifted and set to 0. The same shift was applied to the protein of interest to keep the correct distances between the 2 proteins. The gel expansion factor was applied to all measures before plotting them using GraphPad Prism10. Graphs of the relative average position of the protein of interest according to the tubulin signal were done using the plugin “CentrioleGraph”. For the quantification of the broken phenotype, the number of cells with broken centrioles was manually quantified under the microscope when both or one of the two centrioles per cell were deformed/abnormal or with missing part of the centriole or only blades of microtubules.

#### Expansion factor

For each experiment, the expanded gel is precisely measured with a caliper and the calculated expansion factor is applied in every quantification. Values presented in graphs and scale bars always correspond to “real” values after the application of the expansion factor.

#### siRNA

siRNA efficiency was evaluated manually at the level of the centriole from cells in either G1 (two centrioles) or S/G (four centrioles) phase. The intensity was increased to the maximum, and the signal was monitored. For the measurement of the protein of interest intensity from regular IF or U-ExM, a pixel square with always the same size in between IF measures or in between U-ExM measures was positioned around the centrosome/centriole or in the vicinity to evaluate the background and mean intensity was measured in both regions. Background value was subtracted, and data were plotted as mean intensity values. Data were classified into three categories: signal on the two mature centrioles when all the centrioles were positive for the protein of interest, only one of the two mature centrioles was positive, and completely depleted when the protein of interest was absent from all the mature centrioles.

#### Statistical analysis

Statistical analyses were performed using Excel or Prism10 (GraphPad version 10.2.3 (403), April 21, 2024) and all data are expressed as the mean (average) +/- standard deviation (SD). The comparison of the two groups was performed using an unpaired two-sided Student’s t-test or its non-parametric correspondent, the Mann–Whitney test, if normality was not granted because rejected by the Pearson test. The comparisons of more than two groups were made using one-way ANOVAs for one interaction factor or two-way ANOVAs for several interaction factors followed by multiple comparison tests as indicated in the corresponding Data files to identify all the significant group differences. N indicates independent biological replicates from distinct samples. Every experiment was performed at least three times independently on different biological samples unless specified. No statistical method was used to estimate the sample size. Data are all represented as scatter dot plots with the centerline as mean, except for some percentage quantifications, which are represented as histogram bars. The graphs with error bars indicate SD (+/-) and the significance level is denoted as usual (*P < 0.05, **P < 0.01, ***P < 0.001, ****P < 0.0001).

## Data and software availability

All data are available upon request. The data that support the findings of this study are available as "source data".

## Author Contributions

V.H. and P. G. supervised the present work and wrote the manuscript. L.B performed all the experiments and their analysis with initial help from M.H.L. and S. B.

## Acknowledgments

We thank Juliette Azimzadeh for initial sharing of reagents (plasmid mcherry-CCDC77 and mcherry-WDR67). This work was supported the Swiss State Secretariat for Education, Research and Innovation (SERI) under contract number MB22.00075. This work is supported by the Swiss National Foundation (SNSF) PP00P3_187198 (PG) and 310030_205087 (PG and VH) and by the European Research Council (ERC) ERC StG 715289 ACCENT (PG).

---

**Extended Data Figure 1.**
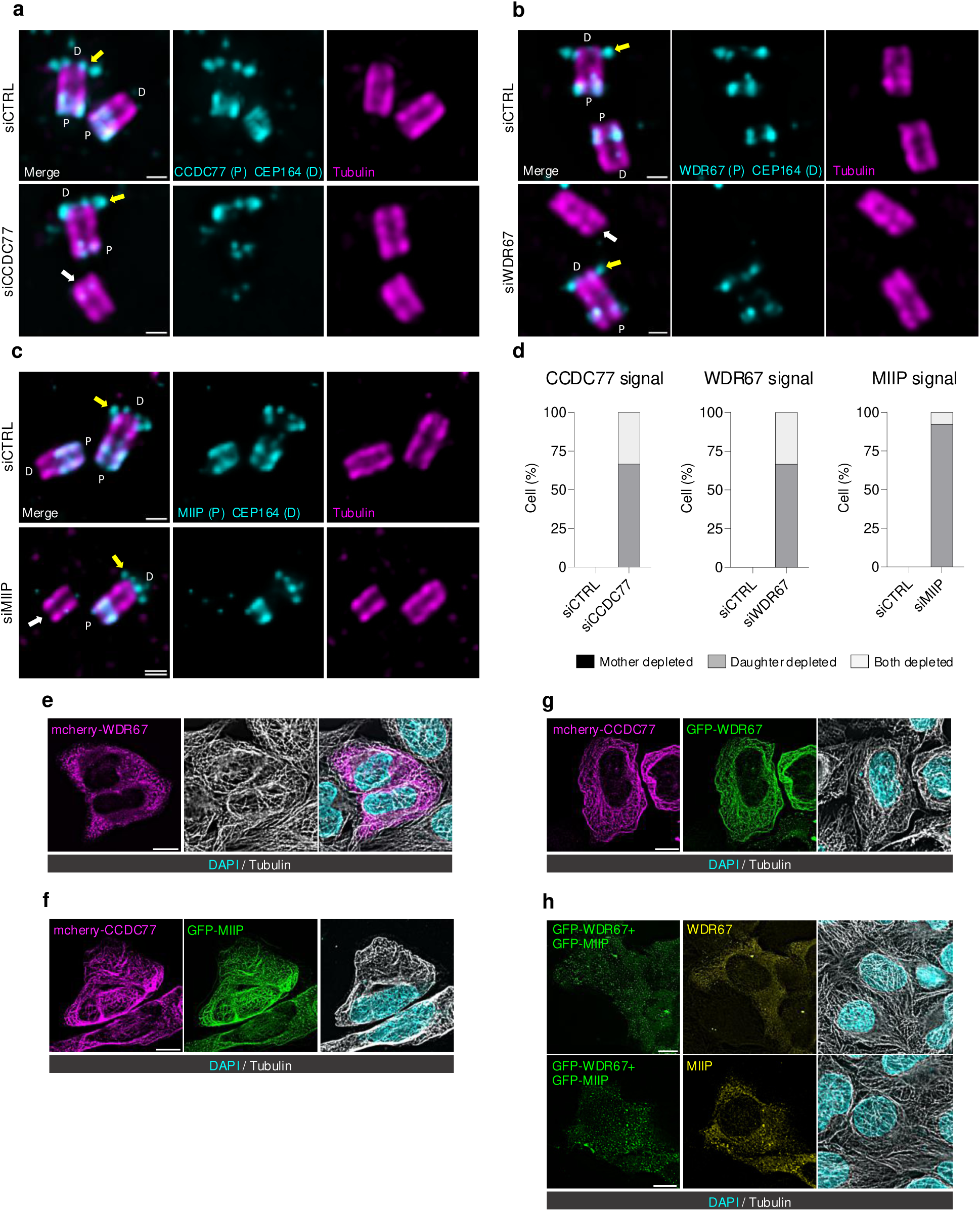
Depletion efficiency and microtubule binding **(a-c)** Widefield images of expanded U2OS centrioles in longitudinal view stained for α/β tubulin (magenta) and CCDC77+CEP164 (cyan) (a), WDR67 + CEP164 (cyan) (b) or MIIP+ CEP164 (c) in the following conditions: siCTRL (a-c) and siCCDC77 (a), siWDR67 (b) or siMIIP (c). P and D stand for proximal and distal part of the centriole respectively. Note that CEP164 serves as a marker for mature centriole (yellow arrow). White arrows point to the signal depletion of the protein of interest for each siRNA treatment. Scale bars = 200 nm. (**d**) Percentage of cells depleted for CCDC77 in siCCDC77, WDR67 in siWDR67, and MIIP in siMIIP on the mother centriole, or the daughter centriole, or both depleted. (**e-g**) Widefield images of U2OS cells transfected with mcherry-WDR67 (e) or with mcherry-CCDC77 and GFP-MIIP (f) or mcherry- CCDC77 and GFP-WDR67 (g) for 24h with doxycycline induction and then after methanol fixation stained for DAPI (cyan) and α/β tubulin (white). Scale bars = 10 μm. (**h**) Widefield images of U2OS cells expressing GFP-WDR67 (green) and GFP-MIIP (green) stained for DAPI (cyan), α/β tubulin (white) and WDR67 (yellow – top panel) or MIIP (yellow – bottom panel). Scale bars = 10 μm. The detailed statistics of all the graphs shown in the figure are included in the Source Data file.

**Extended Data Figure 2.**
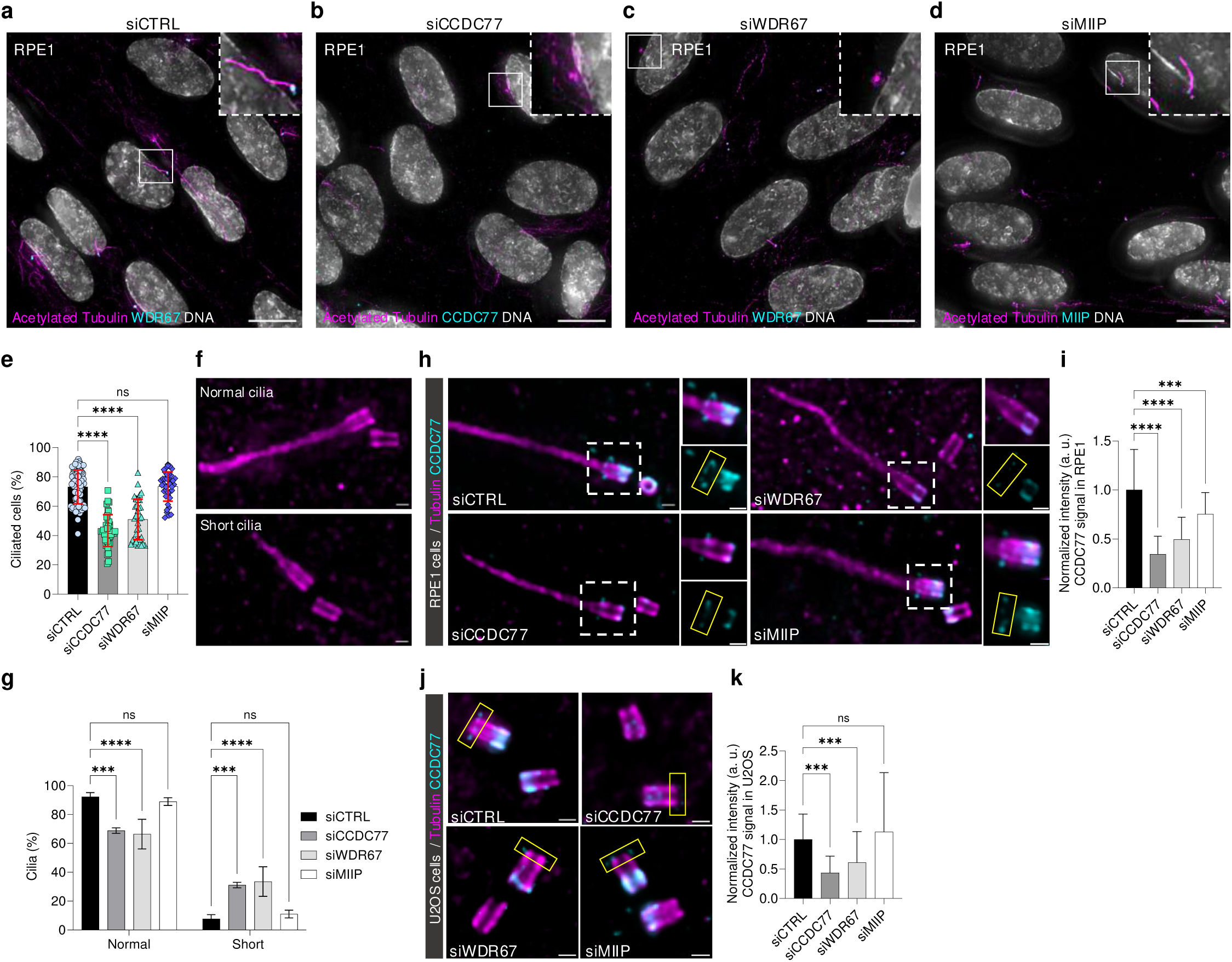
MIIP depletion does not affect ciliogenesis (**a-d**) Widefield images of RPE1 cells ciliated stained for acetylated tubulin (magenta), DAPI (grey) and WDR67 (cyan – a, c), CCDC77 (cyan – b) or MIIP (cyan – d). Scale bars = 10 μm. Dashed-line squares correspond to insets. (**e**) Percentage of RPE1 ciliated cells in the indicated siRNA conditions. (**f**) Widefield images of expanded ciliated RPE1 cells stained for α/β tubulin (magenta). Scale bars = 200 nm. (**g**) Percentage of normal or short cilia in the indicated siRNA conditions. (**h**) Widefield images of expanded ciliated RPE1 cells stained for α/β tubulin (magenta) and CCDC77 (cyan) in different siRNA conditions. Scale bars = 200 nm. Dashed line squares correspond to insets. Yellow squares indicate the distal localization of CCDC77 used for quantification shown in (i). (**i**) Normalized relative intensity of distal CCDC77 signal in the indicated siRNA conditions in RPE1 cells. (**j**) Widefield images of expanded U2OS centrioles in longitudinal view stained for α/β tubulin (magenta) and CCDC77 (cyan) in different siRNA conditions. Scale bars = 200 nm. Yellow squares indicate the distal localization of CCDC77 used for quantification shown in (k). (**k**) Normalized relative intensity of distal CCDC77 signal in the indicated siRNA conditions in U2OS cells. The detailed statistics of all the graphs shown in the figure are included in the Source Data file.

**Extended Data Figure 3.**
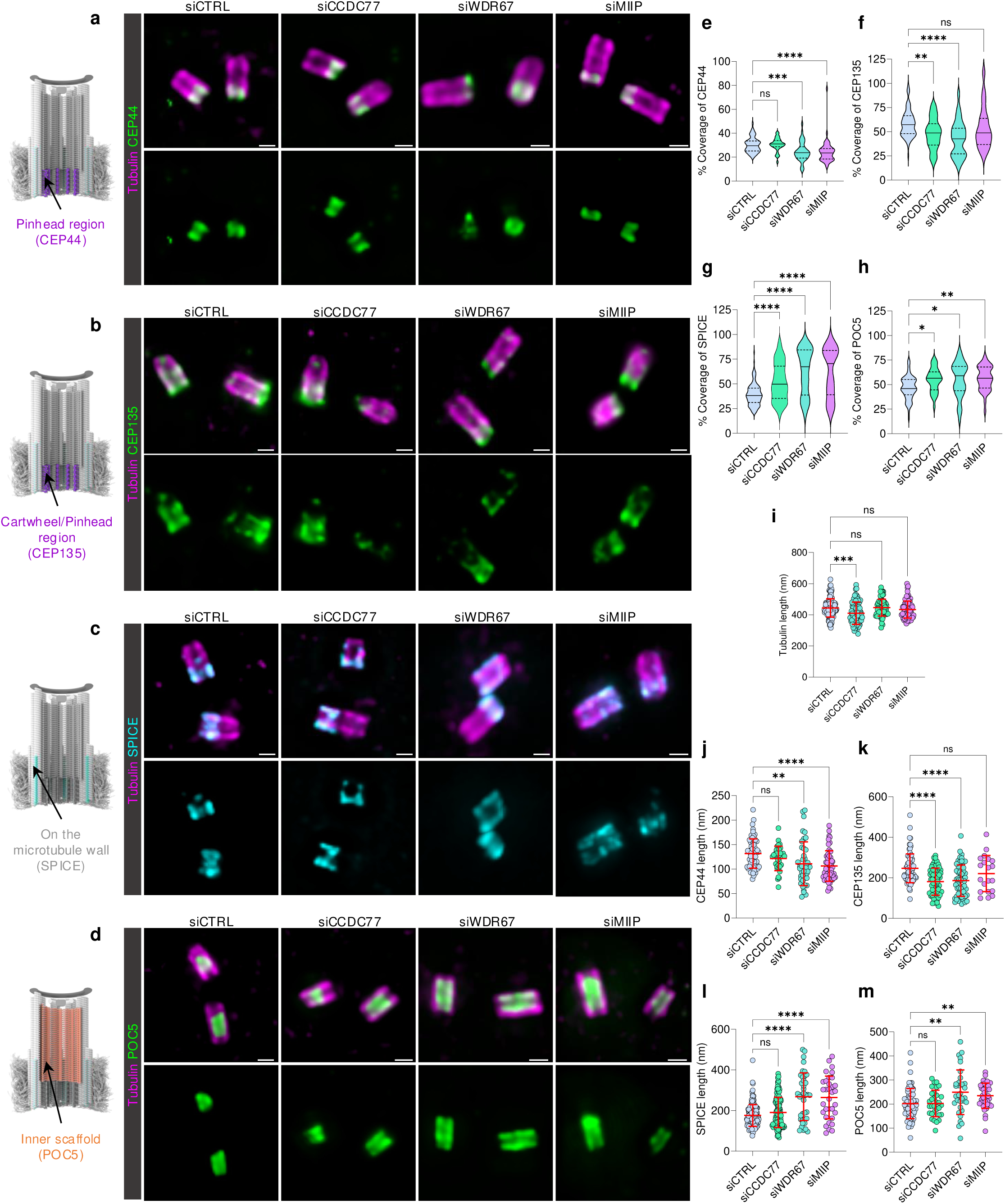
Impact of A-C linker depletion on other centriolar elements. **(a-d)** Widefield images of expanded U2OS centrioles in longitudinal view stained for α/β tubulin (magenta) and CEP44 (a – green), CEP135 (b – green), SPICE (c – cyan) or POC5 (d – green) in the indicated siRNA conditions. Scale bars = 200 nm. Model of a human mature centriole on the left panel displaying the structural element of interest next to the corresponding images. (**e-h**) Coverage of CEP44 (e), CEP135 (f), SPICE protein (g), or POC5 protein (h), expressed as a percentage of the tubulin length in the indicated siRNA conditions. Note that CEP44 and POC5 quantification were performed on daughter centriole only while CEP135 and SPICE on both mother and daughter ones. (**i**) Tubulin length of mature centrioles (mother and daughter) in the indicated siRNA conditions. (**j-m**) Proteins length of CEP44 (j), CEP135 (k), SPICE (l) or POC5 (m) in the indicated siRNA conditions. Note that CEP44 and POC5 quantification were performed on daughter centriole only while CEP135 and SPICE on both mother and daughter ones. The detailed statistics of all the graphs shown in the figure are included in the Source Data file.

**Extended Data Figure 4.**
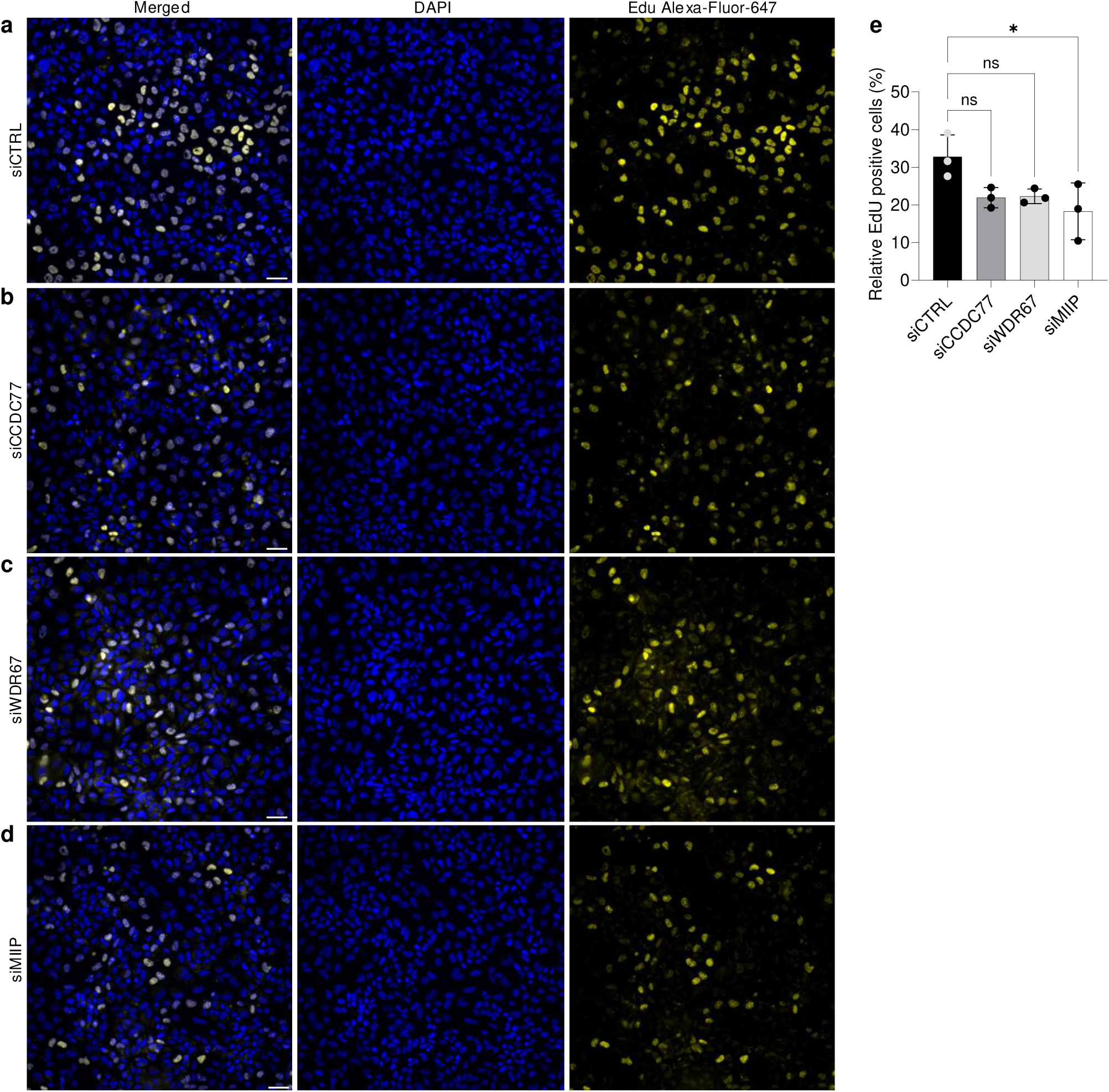
Cell cycle is not impaired upon A-C linker components depletion. (**a-d**) Widefield images of U2OS cells treated with siCTRL (a), siCCDC77 (b), siWDR67 (c) or siMIIP (d) incubated with click-EdU and stained with DAPI (blue) and Edu Alexa-Fluor 647 (yellow). Scale bars = 50 μm. (**e**) Percentage of relative EdU-positive cells in the indicated siRNA conditions. The detailed statistics of all the graphs shown in the figure are included in the Source Data file.

